# Visualizing dendritic characteristics of corticotropin-releasing hormone neurons at single-cell resolution in the whole mouse brain

**DOI:** 10.1101/2020.06.23.168310

**Authors:** Yu Wang, Pu Hu, Qinghong Shan, Chuan Huang, Zhaohuan Huang, Peng Chen, Anan Li, Hui Gong, Jiang-Ning Zhou

**Affiliations:** Chinese Academy of Science Key Laboratory of Brain Function and Diseases, School of Life Sciences, University of Science and Technology of China, Anhui, PR China; Center for Excellence in Brain Science and Intelligence Technology, Chinese Academy of Sciences, Shanghai, 200031, China; Britton Chance Center for Biomedical Photonics, Wuhan National Laboratory for Optoelectronics, MoE Key Laboratory for Biomedical Photonics, School of Engineering Sciences, Huazhong University of Science and Technology, Wuhan 430074, China

## Abstract

Corticotropin-releasing hormone (CRH) is an important neuromodulator with wide distribution in the brain. Here, we screened the CRH-IRES-Cre;Ai32 mouse line to reveal the morphologies of individual CRH neurons throughout the mouse brain by using fluorescence micro-optical sectioning tomography (fMOST) system. Diverse dendritic morphologies and projection fibers were found in various brain regions. Reconstructions showed hypothalamic CRH neurons had the smallest somatic volumes and simplest dendritic branches, and CRH neurons in several regions shared a bipolar morphology. Further investigations in the medial prefrontal cortex unveiled somatic depth-dependent morphologies that exhibited three types of connections and CRH neurons in the anterior parvicellular area of hypothalamus had fewer and smaller Herring bodies whereas in the periventricular area had more and larger Herring bodies that were present within fibers projecting to the third ventricle. Our findings provide the most comprehensive intact morphologies of CRH neurons throughout the mouse brain that is currently available.

## Introduction

Corticotropin-releasing hormone (CRH), a 41–amino acid peptide, is an important neuromodulator with wide distribution in the brain (Spiess et al., 1981). As a neuroendocrine hormone, CRH is abundantly expressed in hypothalamic paraventricular nucleus (PVN) neurons and plays a crucial role in the regulation of the hypothalamic-pituitary-adrenal (HPA) axis (Albeck et al., 1997; Mezey et al., 1984). CRH-expressing neurons are also broadly distributed in other brain regions, including the inferior olivary nucleus, Barrington’s nucleus, pontine tegmentum, cerebral cortex, hippocampus, and central amygdala. Depending on the region-specific somatic locations, CRH neurons participate in various functional activities, such as learning memory, synaptic plasticity, food intake and drug addiction, as well as anxiety-like and depression-like behaviors (Gallopin et al., 2006; Miyata et al., 1999; Puder and Papka, 2001; Sasaki and Sato, 2013).

The anatomy of the brain CRH system has been studied in different mammalian species via immunohistochemistry and radioimmunoassays (Bassett and Foote, 1992; Charlton et al., 1987; Cummings, 1989; Delville et al., 1992; Foote and Cha, 1988; Merchenthaler, 1984; Shu et al., 2015). However, data from these studies have mainly been acquired from histological imaging and through manual reconstruction and counting of labeled neurons, which is time-consuming, limits further systematic analysis, and can introduce biases and/or artifacts. Recently, genetically modified mouse models have been developed to identify the whole-brain distributions of CRH neurons (Itoi et al., 2014; Kono et al., 2017; Peng et al., 2017; Wamsteeker Cusulin et al., 2013), which has significantly advanced our understanding of the morphological features of CRH neurons in the rodent brain (De Francesco et al., 2015; Garcia et al., 2016; Huang et al., 2013; Nguyen et al., 2016). Advances in whole-brain optical imaging techniques, such as fluorescence micro-optical sectioning tomography (fMOST) (Gong et al., 2016; Gong et al., 2013; Ragan et al., 2012; Zheng et al., 2013), have made it feasible to further quantify cellular distributions and to morphologically reconstruct cells at the whole-brain level. The precision of imaging via fMOST can reveal complex fiber orientations, and can even distinguish individual dendrites. Such quantitative three-dimensional (3D) neuronal morphologies obtained at a brain-wide scale can provide highly accurate arborization details and comprehensive mapping of CRH neuronal connections throughout the brain.

Although the whole-brain expression patterns of CRH have been qualitatively analyzed (Alon et al., 2009; Kono et al., 2017; Peng et al., 2017), high-resolution reconstruction of the full morphologies (including both somata and dendrites) of CRH neurons at the single-neuron level has rarely been performed. Since neuronal morphology is considered to be one of the most defining features to distinguish among neuronal types and network connectivities, characterizing single-neuron morphologies may provide key information on how neuronal information and signals are transmitted within the local networks. Furthermore, analysis of detailed morphological information (including data sets of somatic locations, as well as dendritic and axonal morphological features) of diverse CRH neurons may facilitate in better classifying CRH neuronal types and in revealing their local connectivity. For example, a recent study employed fMOST to investigate CRH distributions in the mouse brain, enabling quantitative analysis of whole-brain CRH somata (Peng et al., 2017) that provides us with substantial quantitative information on brain CRH networks. However, at present, the morphological details of individual CRH neuronal fibers at the whole-brain level remain poorly understood. In the present study, we constructed a comprehensive whole-brain map of genetically labeled CRH neurons in the mouse brain, which provides dendritic distribution patterns of single CRH neurons. Reconstructions and further analysis showed that heterogeneous CRH interneurons in the mPFC form layer-dependent dendritic-dendritic and dendritic-somatic connections; and that there was a target-oriented distribution of large vesicles (Herring bodies) within the processes of hypothalamic CRH neurons. This work provides a comprehensive description of the whole-brain CRH neuronal distribution pattern and, more importantly, dendritic morphological features of CRH neurons in the mouse brain.

## Results

### Comparison of morphological features of fluorescent-labeled CRH neurons in three fluorescent-reporter mouse lines

By crossing CRH-IRES-Cre mice with Ai6, Ai14, and Ai32 reporter mice, in which the cassette containing ZsGreen1, td-Tomato, or CHR2-EYFP was expressed in a Cre-dependent manner, we obtained CRH-IRES-Cre;Ai6, CRH-IRES-Cre;Ai14, and CRH-IRES-Cre;Ai32 mice, respectively (Fig. S1A). Then, we compared the distributions and morphologies of fluorescent-labeled CRH neurons in several brain regions, including the olfactory bulb (OB) (Fig. 1A–C), cortex (Fig. 1D–F), PVN (Fig. 1G–I), bed nucleus of the stria terminalis (BST) (Fig. S1B–D) and central nucleus of the amygdala (CeA) (Fig. S1E– G).

**Figure 1.**
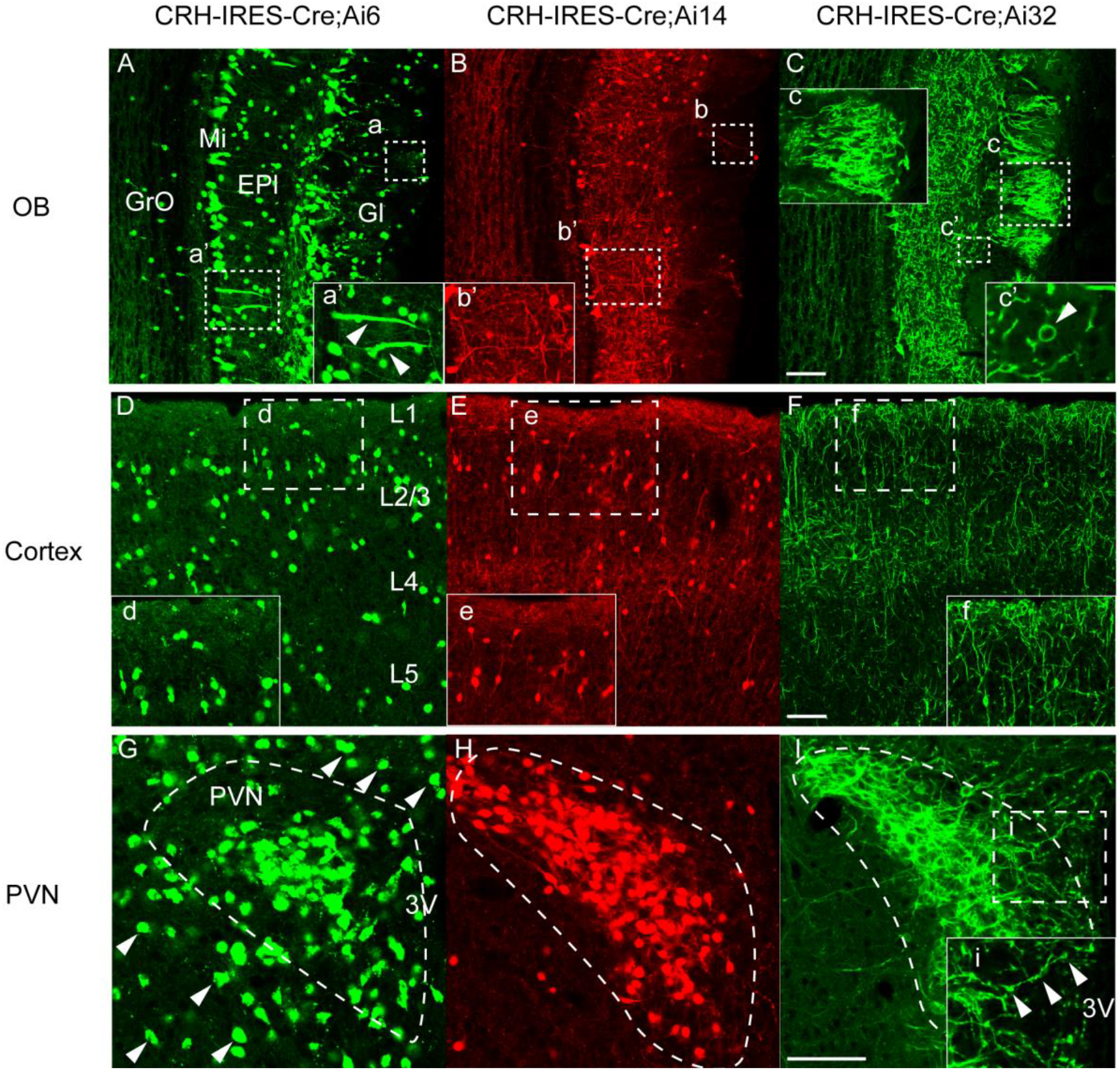
Comparison of the morphological features of CRH neurons in three fluorescent-reporter mouse lines. Comparisons of the distributions and morphologies of fluorescent-labeled CRH neurons in several brain regions of CRH-IRES-Cre;Ai6, CRH-IRES-Cre;Ai14, and CRH-IRES-Cre;Ai32 mice in the OB (A–C), cortex (D–F), and PVN (G–I). (A–C, a–c) The dotted boxes show ZsGreen1 (a), td-Tomato (b) and EYFP (c) labeling of synaptic globular structures in the Gl of the three mouse lines. (A–B, a’–b’) The dotted boxes and magnified images show ZsGreen1 (a’) and td-Tomato (b’) labeling of cell bodies and dendrites (indicated by the arrowheads) in the Mi and EPl of CRH-IRES-Cre;Ai6 and CRH-IRES-Cre;Ai14 mice. (C, c’) The dotted box and magnified image show EYFP-labeled cell body (indicated by the arrowhead) and dendrites in a CRH-IRES-Cre;Ai32 mouse. (D–F, d–f) The dotted boxes and magnified images show ZsGreen1 (d), td-Tomato (e), and EYFP (f) labeling of cell bodies and dendrites in layer 1 and layer 2/3 of the three mouse lines. (G) ZsGreen1-labeled cell bodies in (dotted-curved box) and outside (arrowheads) of the PVN of CRH-IRES-Cre;Ai6 mice. (H) Td-Tomato-labeled cell bodies in the PVN (dotted-curved box) of CRH-IRES-Cre;Ai14 mice. (I and I, i) EYFP-labeled cells (I, dotted-curved box) and fibers (i, dotted box and magnified image) in the PVN of CRH-IRES-Cre;Ai32 mice. Scale bars = 100 μm.

In the OB, the transgenic fluorescent proteins were mainly distributed in the glomerular layer (Gl), external plexiform layer (EPl), and the mitral cell layer (Mi) in all three mouse lines. CRH-IRES-Cre;Ai6 mice showed the brightest and largest number of fluorescent-labeled cells in all these layers (Fig. 1A); in particular, more fluorescent cells were labeled in the granule cell layer (GrO) in this mouse line compared to those in CRH-IRES-Cre;Ai14 and CRH-IRES-Cre;Ai32 mice. However, fluorescent-labeled neuronal fibers were short and their fluorescent distributions were not uniform; for example, the dendrites close to the cell bodies of mitral cells were strongly labeled (Fig. 1A, a’, indicated by the arrowheads), but the branches extending to the Gl were not clear (Fig. 1A, a, indicated by the dotted box). CRH-IRES-Cre;Ai14 mice were also labeled with bright cell bodies and dense fibers were labeled in the EPl (Fig. 1B, b’), but only a few dendritic structures were labeled in the Gl (Fig. 1B, b, indicated by the dotted box). By contrast, in each layer of CRH-IRES-Cre;Ai32 mice, the fluorescence distributed in cell bodies and fibers exhibited a uniform brightness (Fig. 1C), and the somata in these sections were organized in a ring-like structure (Fig. 1C, c’, indicated by the arrowhead). Unlike the former two mouse lines, the Gl showed a bushy spherical structure (Fig. 1C, c) that was comprised of mitral cells and/or peribulbar cells.

In the cortex, fluorescent-labeled cells were found in each layer in CRH-IRES-Cre;Ai6 mice (Fig. 1D). The cell bodies were strongly labeled, while fibers were rarely seen (Fig. 1D, d). The numbers of labeled cells in the cortices of CRH-IRES-Cre;Ai14 and CRH-IRES-Cre;Ai32 mice were less than those in CRH-IRES-Cre;Ai6 mice, and the cells were mainly distributed in layer 2/3 (Fig. 1E–F). Neurons in CRH-IRES-Cre;Ai14 mice also showed clearer and brighter cell bodies (Fig. 1E, e), whereas more fibers were labeled (Fig. 1F) in CRH-IRES-Cre;Ai32 mice, especially in terms of a dense distribution in the first layer (Fig. 1F, f).

The outlines of nuclei were clearly visible in the fluorescent labeling of CRH neurons in the PVN (Fig. 1G–I, indicated by the dotted line), BST (Fig. S1B–D, indicated by the dotted box) and CeA (Fig. S1E–G, indicated by the dotted box) of the three mouse lines. Similarly, CRH-IRES-Cre;Ai6 mice showed the highest number of labeled neurons with brighter cell bodies in these three regions (Fig. 1G; Fig. S1B, a and Fig. S1E, d). In CRH-IRES-Cre;Ai14 mice, distinguishable cell bodies and dense fibers were labeled in the BST (Fig. S1C, b) and CeA (Fig. S1F, e), while only the cell bodies were clearly seen in the PVN (Fig. 1H, indicated by the dotted line). By contrast, CRH-IRES-Cre;Ai32 mice showed dense fibers in all of these regions, and the fluorescent signals of the cell bodies were distinguishable (Fig. 1I, Fig. S1D, c and Fig. S1G, f); especially in the PVN, neuronal fibers extending to the lateral and third ventricle (Fig. 1I, i, fibers indicated by the arrowheads) were visible, and there was a uniform fluorescent intensity distributed in the nearby cell bodies and fibers.

In summary, among the three reporter mouse lines, CRH-IRES-Cre;Ai6 and CRH-IRES-Cre;Ai14 mice showed clearer and brighter cell bodies of CRH neurons. CRH-IRES-Cre;Ai6 mice had the largest number of labeled CRH cells in each brain region, but almost no neuronal fibers were visible. CRH-IRES-Cre;Ai14 mice showed clear but incomplete fibers. Only CRH-IRES-Cre;Ai32 mice showed the most complete fibrous structures, especially in terms of distributions in neuronal terminals (e.g., the bushy spherical structures in the glomerular layer of the OB; the extended fibers in the cortex and PVN); Regardless of its weaknesses on distinguishing single cell bodies, the fluorescent distributions in were uniform, which is conducive to the adjustment of exposure and the collection of complete morphologies of neurons during imaging.

### Whole-brain distributions of CRH neurons at high resolution in the CRH-IRES-Cre;Ai32 mouse line

Since the single-cell morphology of CRH neurons was most clearly visible in CRH-IRES-Cre;Ai32 mice, we used this mouse line to image EYFP-labeled CRH neurons throughout the brain at a resolution of 0.2 × 0.2 × 1.0 μm via an fMOST system. First, 100-μm down-sampled coronal projection sections (Fig. 2A) were provided to show the overall distributions of CRH neurons in various brain regions. EYFP-labeled cells were distributed in many regions that has not been reported such as in vascular organ of the lamina terminalis (VOLT), ventromedial preoptic nucleus (VMPO), caudate putamen (CPu), bed nucleus of the anterior commissure (BAC), triangular septal nucleus (TS), suprachiasmatic nucleus (SCN), Kölliker-Fuse nucleus (KF) and nucleus X (X) (Fig. 2A). Furthermore, bundles of CRH projection fibers were visible in accumbens nucleus, shell (AcbSh), interstitial nucleus of the posterior limb of the anterior commissure (IPAC), anterior commissure, posterior (acp), corpus callosum (cc), and inferior cerebellar peduncle (icp) (Fig. 2A, indicated by arrowheads and Fig. 2B). The movies of serial sections showed that the fibers in AcbSh, IPAC, and acp were projections from neurons in the OB (Movie S1) and fibers in icp were projections from IO CRH neurons (Movie S2). Moreover, we found novel populations of CRH-positive neurons in some brain regions, such as a sparsely distributed group in the CPu (Fig. 2C) that had dendrites that were radially distributed (with the maximum radius from the terminals to the somata being 40–70 μm). Neurons gathered in the BAC (Fig. 2D) had round cell bodies and two short processes. The average number of CRH-positive neurons found in the SCN was 45.67±0.88, and nearly every neuron had two thick primary dendrites with few branches (Fig. 2E). Neurons in the dorsal cochlear nucleus (DC) (Fig. 2F) had dense apical dendrites distributed in the superficial glial zone, and chandelier cells with apical dendrites vertically distributed were labeled in the cerebellum (Fig. 2G). A cluster of swollen structures (Fig. 2H, indicated by arrowheads) presenting transparent smooth surfaces was visible around the third ventricle (3V) and always extended to the 3V border. Densely labeled vascular-like structures and terminals of CRH neurons were found in VOLT (Fig.2 I) and ME (Fig. 2J). The distributions and morphologies of CRH neurons in other brain regions are shown in Fig. S2.

**Figure 2.**
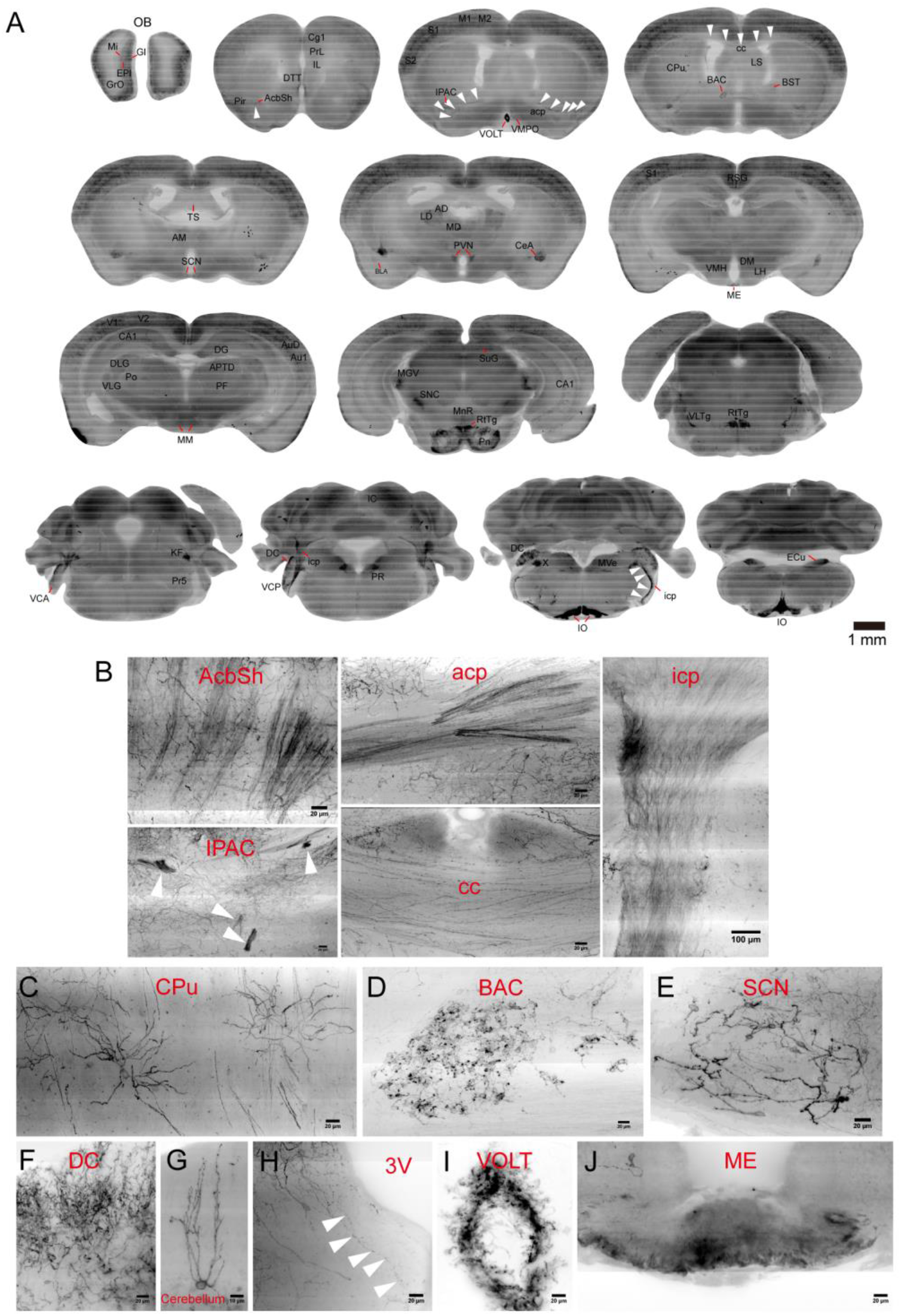
Whole-brain distributions of CRH neurons at high resolution in the CRH-IRES-Cre;Ai32 mouse line. (A) Whole-brain distributions of EYFP-labeled CRH neurons, using the fMOST system, contained in 100-μm coronal-projected sections showing the general distributions of EYFP-labeled CRH neurons in different brain regions. (B) EYFP-labeled clusters of fibers projections from CRH neurons in the AcbSh, IPAC, acp, cc, and icp. (C) EYFP-labeled single neurons in the CPu. (D) Clustered EYFP-labeled neurons distributed in the BAC. (E) EYFP-labeled neurons in the SCN. (F) Dense EYFP-labeled dendrites from neurons in the DC. (G) EYFP-labeled single neuron in the cerebellum. (H) EYFP-labeled fibers and swollen structures around the 3V. (I) EYFP-labeled vascular like structures in the VOLT. (J) EYFP-labeled terminals of PVN CRH neurons in the ME.

### Three-dimensional distributions and single-cell reconstructions of CRH neurons in several brain regions

We reconstructed EYFP-labeled CRH neurons in several brain regions (Fig. 3A–H), including the OB (Fig. 3 A), dorsal part of lateral septal nucleus (LSD) (Fig. 3B), BST (Fig. 3C), CeA (Fig. 3D), VMPO (Fig. 3E), hippocampus (Hip) (Fig. 3F), SCN (Fig. 3G), and DC (Fig. 3H). We found that the reconstructed neurons in several brain regions (e.g., mPFC, BST, VMPO, anterior parvicellular part of paraventricular hypothalamic nucleus (PaAp), periventricular hypothalamic nucleus (Pe), and SCN) shared similar morphological characteristics consistent with bipolar neurons (Fig. 3I). CRH neurons in the LSD had the largest average volume of somata (1632±159.6 μm^3^) (Fig. 3J) and the longest dendritic length (1.9±0.2 mm) (Fig. 3K). Dendritic length significantly increased as a function of somatic volume (R2=0.597, P=0.0032) (Fig. 3N). CRH neurons in the VMPO also had a larger cell bodies (1172±228.1 μm^3^) (Fig. 3J), but the number of dendritic branches (15.2±3.7) (Fig.3 L) and dendritic length (1.1±0.1 mm) (Fig. 3K) were less than those of neurons in the LSD. Dendritic length was also positively correlated with somatic volume in the VMPO (Fig. 3O). Sholl analysis showed that neurons in the LSD had the largest maximum number of intersections, while the VMPO had the least maximum number of intersections. For all of these regions, the maximum numbers of intersections were located at radial distances of 50–100 μm from the somata (Fig. 3M). The more complex dendrites of CRH neurons in the LSD, compared to those in other areas, suggested that CRH neurons in the LSD may receive comparatively more inputs. Most of the dendritic morphologies of CRH neurons in the hippocampus exhibited a similar pattern of an umbrella shape of upward dendrites (Fig. 3F). CRH neurons in the SCN were scattered throughout the nucleus and the dendrites were interlaced with one another (Fig. 3G). We next compared the parameters of all reconstructed neurons (Table S1) in different brain regions and found that CRH neurons in hypothalamic regions—including the PaAp (640.1±60.4 μm^3^), Pe (951.2±108.3 μm^3^), and SCN (636.0±55.4 μm^3^)—had smaller somatic volumes (Fig. 5J and Table S1). Similarly, there were also shorter dendritic lengths of CRH neurons in the PaAp (0.5±0.02 mm), Pe (0.5±0.06 mm), and SCN (0.6±0.05 mm) (Fig. 3K and Table S1). The simpler morphologies of hypothalamic CRH neurons may be related to their endocrine and other conserved functions.

**Figure 3.**
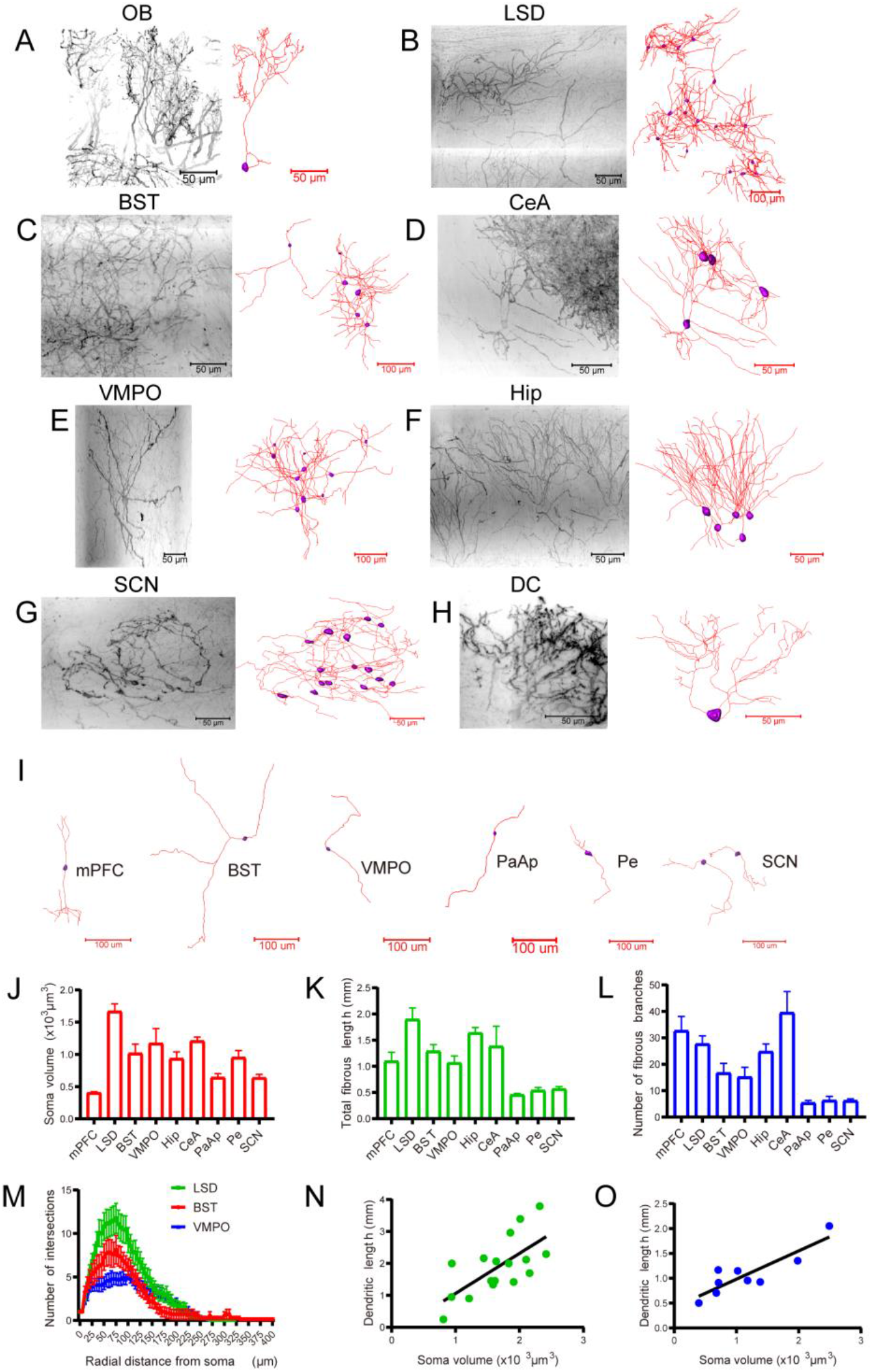
Three-dimensional distributions and single-cell reconstructions of CRH neurons in several brain regions. (A–H) Original images (left half) and reconstructions (right half) of CRH neurons in different brain regions, including the OB (A), LSD (B), BNST (C), CeA (D), VMPO (E), Hip (F), SCN (G), and DC (H). The original images were inverted into grayscale images. The reconstructed somata and dendrites of neurons are indicated by purple bodies and red lines, respectively. (I) Typical reconstructed neurons show the common bipolar morphology found in different brain regions, including the mPFC, BNST, VMPO, PaAp, Pe, and SCN. (J) Somatic volumes of CRH neurons in different brain regions [one-way ANOVA, P < 0.0001, F (8, 125) = 23.24]. (K) Total fibrous lengths of CRH neurons in different brain regions [one-way ANOVA, P < 0.0001, F (8, 95) = 12.26]. (L) Numbers of fibrous branches of CRH neurons in different brain regions [one-way ANOVA, P < 0.0001, F (8, 101) = 11.53]. (M) Sholl analysis of dendrites of neurons in the LSD, BNST, and VMPO illustrate changes in the mean number of intersections with increasing radial distance from the soma. (N) Correlation between dendritic length and somatic volume in LSD neurons (Pearson’s correlation coefficient r = 0.68, P = 0.0029). (O) Correlation between dendritic length and somatic volume in VMPO neurons (Pearson’s correlation coefficient r = 0.88, P = 0.0019).

### Multiple morphological types of CRH neurons form distinct dendritic connections in the medial prefrontal cortex (mPFC)

Recent studies have shown that CRH neurons in the mPFC play a critical role in higher cognitive functions (Chen et al., 2020; Hupalo et al., 2019b). In the prelimbic cortex (PrL) within the mPFC, we reconstructed the entire somata (Fig. 4A, purple bodies) and dendrites (Fig. 4A, color lines) of EYFP-labeled CRH neurons within a column that had a volume of 350 × 500 × 500 μm. The cell bodies of these neurons were mostly distributed within layers 2–4 and most of their dendrites were vertically distributed. There were dendritic branches in both the upper and lower parts of the somata. The apical dendrites that branched in the first layer formed a dense dendritic network, and most of them reached the pia mater (Fig. 4A, reconstructed fibers indicated in layer 1); furthermore, the basal dendrites extended and branched into layer 4 at a distance of approximately 500 μm from the cortical surface (Fig. 4A, reconstructed fibers indicated in layer 4). Individual reconstructed neurons were classified according to the distances (50–100, 100–150, 150–200, 200–250, and >250 μm) between their somata and the surface of the cortex (Fig. 4B), and the percentages of neurons in these categories were 19%, 31%, 35%, 10%, and 5%, respectively (Fig. 4B, indicated in the pie chart). We found that there was significant correlation between the somatic depth (distance from the cortical surface) and both the total dendritic length (Fig. 4C, upper half, r=0.4440) and total Euclidean distance (the straight-line distance from the soma to the given point of the dendrite) (Fig. 4C, bottom half, r=0.5399). Sholl analyses showed that the number of intersections with a radial distance from the soma being less than 50 μm was larger in neurons with a somatic depth of 50–100 μm than that of neurons with a somatic depth of 100–150 μm; in contrast, the number of intersections with a radial distance from the soma being more than 50 μm was smaller and ended at approximately 100 μm of the radial distance from the soma in neurons with a somatic depth of 50–100 μm. Interesting, the maximum numbers of intersections were similar between these two types of neurons (Fig. 4D). We also found that the total dendritic length (Fig. 4E, left half) and the total Euclidean distance (Fig. 4E, right half) of neurons with a somatic depth of less than 100 μm were significantly smaller than those with somatic depths of 100–150 μm (P=0.0285) and more than 150 μm (P=0.0005), while there were no significant differences in the total number of dendritic branches or the total number of dendritic terminal points (Fig. S3I). The average dendritic lengths of neurons at somatic depths of less than 100 μm, 100–150 μm, and more than 100–150 μm were 0.82±0.26, 1.33±0.54, and 1.48±0.68 mm, respectively; furthermore, their total Euclidean distances were 14.65±5.19, 38.77±25.49, and 48.67±30.38 mm, and their total numbers of dendritic branches were 23.23±8.65, 32.86±20.28, 33.32±19.44, respectively.

**Figure 4.**
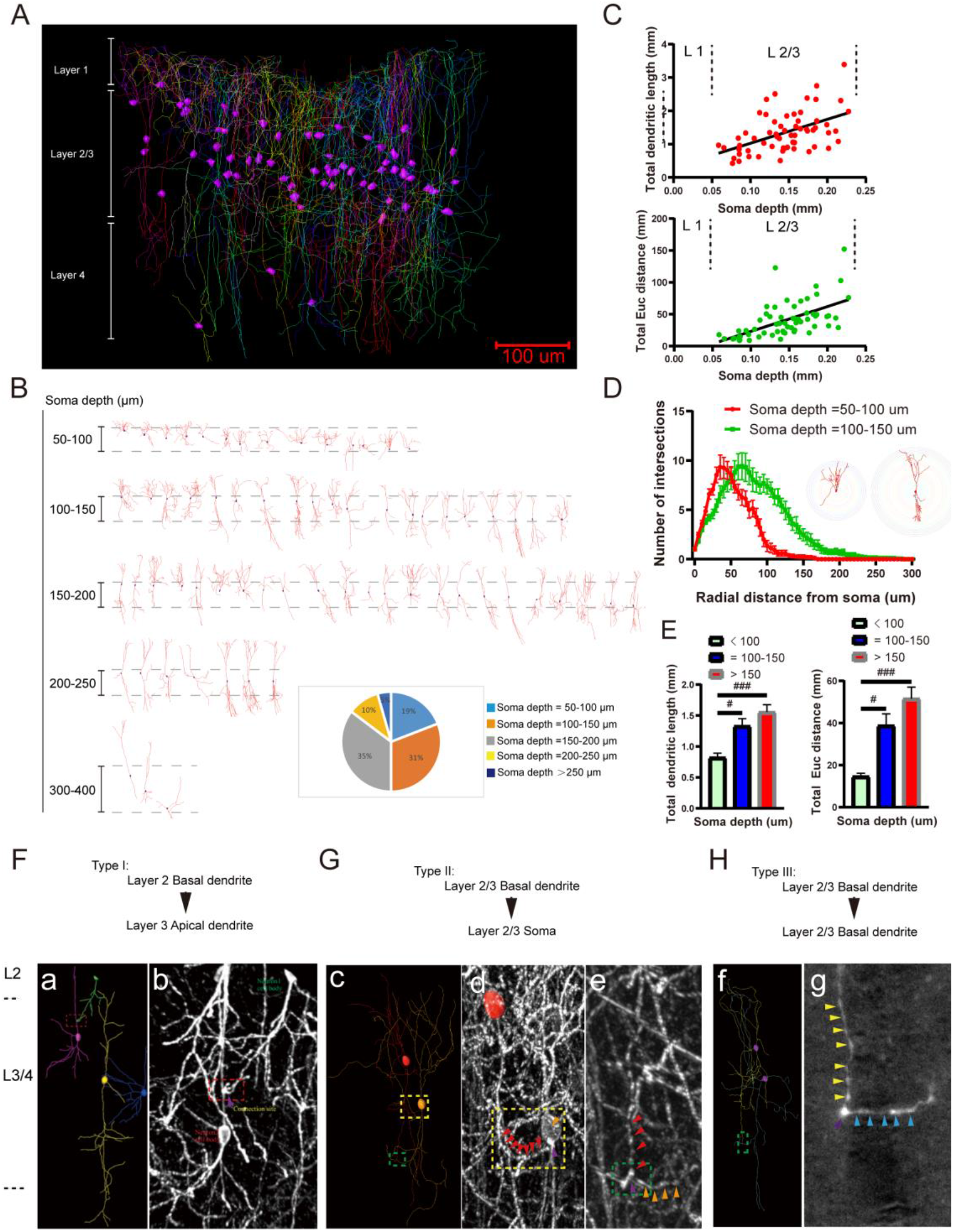
Multiple morphological types of CRH neurons form putative connections in the mPFC. (A) Overview of reconstructed CRH neurons including somata (indicated by the purple bodies) and fibers (indicated by the colored lines) from layers 1–4 in a column with a volume of 350 × 500 × 500 μm in a PrL subregion within the mPFC. (B) Representation of 67 reconstructed somata and dendrites of CRH neurons arranged along their somatic depths with respect to the pial surface; the pie chart at the bottom right shows the percentages of neurons distributed at different somatic depths. (C) Upper half: Correlation between total dendritic length and somatic depth in PrL neurons (Pearson’s correlation coefficient r = 0.51, P<0.0001); Lower half: Correlation between total Euclidean distance and somatic depth in PrL neurons (Pearson’s correlation coefficient r = 0.59, P<0.0001). (D) Sholl analysis of the dendrites of neurons with somatic depths = 50–100 μm and 100–150 μm illustrating changes in the mean number of intersections with increasing radial distance from the soma; Inset images show exemplary intersections on two typical neurons (left: somatic depth = 50–100 μm, right: somatic depth = 100–150 μm) with different somatic depths. (E) Left: The total dendritic length of CRH neurons with somatic depths < 100 μm showed differences compared with those with somatic depths = 100–150 μm (one-way ANOVA, multiple comparisons, P = 0.0285) and soma depths >150 μm (one-way ANOVA, multiple comparisons, P = 0.0005). Right: The total Euclidean distance of CRH neurons with a somatic depth of < 100 μm showed differences compared with those with soma depths = 100–150 μm (one-way ANOVA, multiple comparisons, P = 0.0213) and somatic depths > 150 μm (one-way ANOVA, multiple comparisons, P = 0.0001). (F–H) Three types of connections between CRH neurons in the mPFC. (F) a (reconstructed neurons) and b (original image) show the type-I connection (the connection site is indicated by the purple arrowhead in the red dotted box). (G) The yellow-dotted boxes in c (reconstructed neurons) and d (original image) show the type-II connection (red arrowheads indicate a branch of basal dendrites of one neuron and the orange arrowhead indicates the cell body of another neuron; the connection site is indicated by the purple arrowhead). Green-dotted squares in c (reconstructed neurons) and e (original image) showed the type-III connection (red arrowheads indicated a branch of basal dendrites of one neuron and the orange arrowheads indicate a branch of basal dendrites of another neuron; the connection site is indicated by the purple arrowhead). (H) Green-dotted squares in f (reconstructed neurons) and g (original image) also showed the type-III connection, (yellow arrowheads indicate a branch of basal dendrites of one neuron and the blue arrowheads indicate a branch of basal dendrites of another neuron; the connection site is indicated by the purple arrowhead).

To investigate local CRH-CRH connection patterns within the cortex, we divided CRH-CRH connections into three types (Fig. 4F–H). Type I consisted of basal-to-apical connections. Here, somata in layer 2 sent dendrites downward (the green cell of fig.4 F-a) that contacted with the upward dendrites (the purple cell of Fig. 4F, a) from the somata in layer 3 (as shown in the red dotted box of Fig. 4F, a and b). Type II consisted of basal-to-somatic connections (as shown in the yellow dotted box of Fig. 4G, c and d). In layer 2–3, a soma in the upper layer sent dendrites downward (as shown in Fig. 4G, c, red cell), and the end of one branch (Fig.4 G-d red arrows) was in contact with an adjacent lower cell body (Fig. 4G, c, orange cell; Fig. 4G, d, orange arrows). Type III consisted of basal-to-basal connections (Fig. 4G, c and e; Fig. 4H, f, green dotted box). Two cell bodies in layer 2–3 sent dendrites downward, and the end of one branch (Fig. 4G, e, red arrows; Fig. 4H, g, yellow arrows) from the upper soma and the branch (Fig. 4G, e, orange arrows; Fig. 4H, g, blue arrows) from the lower soma formed a connection. A common feature of the three types of connections was that the fluorescent intensity increased at the contact point, indicating a possible connection of structures (Fig. 4F, b, G, d and e, and H, g, purple arrows). The connections of type II and type III are demonstrated in movie S3.

We next performed immunofluorescent staining to determine the specificity of EYFP-labeled neurons in CRH-IRES-Cre;Ai32 mice. The results showed that most of the EYFP-labeled neurons in the mPFC were co-labeled with CRH antibodies (Fig. S3 A–C, indicated by white arrow). Interestingly, in adult hybrid mice, EYFP-labeled pyramidal neurons were visible in layer 3 or layer 5 of the cortex (Fig. S3F), but there were no EYFP-labeled pyramidal neurons on the 21st day after birth (Fig. S3E). These pyramidal neurons did not co-exist with CRH antibodies (Fig. S3G), including their dendrites and spines (Fig. S3, a, indicated by arrowheads). We also observed that some EYFP-labeled neurite swellings in layer 1 were also labelled with CRH antibodies (Fig. S3 b, indicated by arrowheads).

### Reconstructions and morphological features of CRH neurons in the PaAp and Pe

Hypothalamic neuroendocrine CRH neurons play an important role in stress responses, but neurons within different subregions require more detailed morphological analysis. We chose EYFP-labeled neurons in the PaAP and Pe to reconstruct their somata and processes (Fig. 5A and C; Fig. S4D and E). There was a noteworthy co-localization pattern (Fig. S4A–C) for the EYFP-labeled signals and CRH antibodies in the PaAP. There were vesicular fluorescent labels (Herring bodies) (Fig. 5B and D, grey reconstructed structures) on the neurites of neurons in both the PaAP and Pe, and they also co-labeled with CRH antibodies (Fig. 5E, indicated by arrowheads). In terms of their 3D patterns, the somata of some neurons (Fig. 5A–B, purple reconstructed cell bodies) distributed in the PaAP sent out fibers rostrally (Fig. 5A–B, red lines), and there were spaced and small Herring bodies (Fig. 5B, b–d, grey bodies) on these fibers. An example of these reconstructed cells is shown (Fig. 5B) according to the primary branches and number of Herring bodies, and there were four distribution patterns of these neurons. Pattern 1 (Fig. 5B, a) consisted of cells that had two primary branches with the shortest fiber length and no Herring bodies. Pattern 2 (Fig. 5B, b) consisted of cells that had two primary branches with similar fiber lengths at both ends of the somata and were distributed almost vertically, and the fibers extending rostrally had Herring bodies. Pattern 3 (Fig. 5B, c) consisted of cells with both ends of the fibers having Herring bodies and the one fiber branch that was distributed horizontally was longer and extended rostrally, whereas the other short branch was distributed vertically. Finally, pattern 4 (Fig. 5B, d) consisted of fibers of multipolar cells extending rostrally having Herring bodies and being distributed horizontally. The locations of the reconstructed somata in the PaAP are shown in Fig. S4D (purple bodies indicated by red circles).

**Figure 5.**
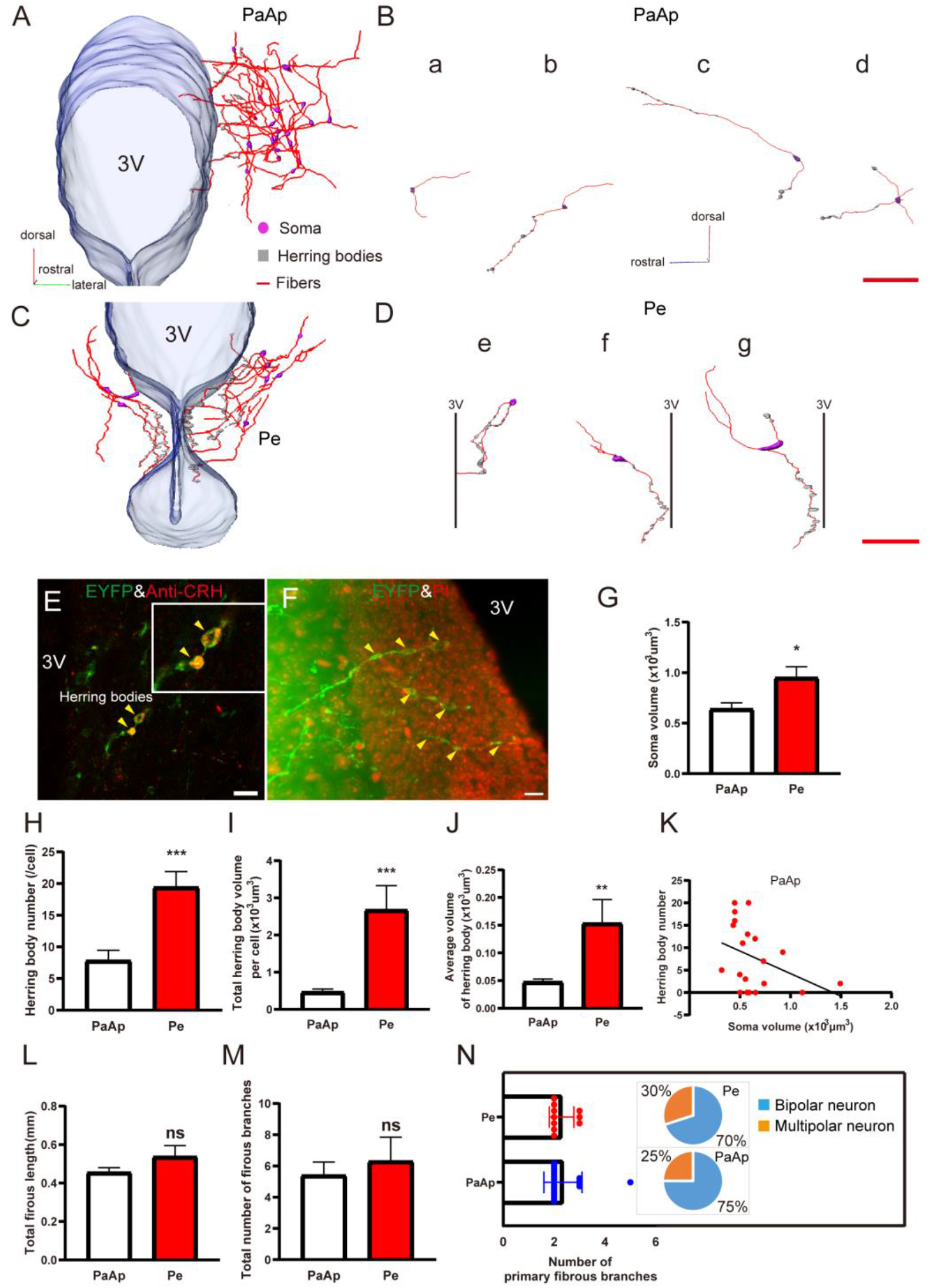
Reconstructions and morphological features of CRH Neurons in the PaAp and Pe. (A) Three-dimensional distribution of reconstructed whole morphologies of CRH neurons in the PaAp. The somata are indicated by the purple bodies, the grey bodies indicate the Herring bodies, and the red lines indicate the neural fibers; the transparent structure on the left represents the reconstructed 3V. (B) Four typical reconstructed neurons in the PaAp. B-a shows a bipolar neuron with short fibers and no Herring bodies; B-b shows a bipolar neuron with Herring bodies located on the fiber at one end of the cell body. B-c shows a bipolar neuron with Herring bodies located on fibers at both ends of the cell body; B-d shows a multipolar neuron with Herring bodies located on the neural fibers (scale bar = 100 μm). (C) Three-dimensional distribution of reconstructed whole morphologies of CRH neurons in the Pe. The somata are indicated by purple bodies, the grey bodies indicate the Herring bodies, and the red lines indicate the fibers; the transparent structure in the center represents the reconstructed 3V. (D) Three individual reconstructed neurons in the Pe. D-e shows a bipolar neuron with both ends of fibers extending to the 3V and with Herring bodies located on fibers at both ends of the cell body; D-f shows a bipolar neuron with one end of the fiber extending to the 3V and with Herring bodies located on the fiber at one end of the cell body; D-g show a multipolar neuron with one fiber extending to the 3V and with Herring bodies mostly located on the fiber at one end of the cell body (scale bar = 100 μm). (E) EYFP-labeled Herring bodies were CRH-antibody-positive (indicated by yellow arrowheads, inset image is a magnified image). (F) The fiber ends with Herring bodies located (indicated by yellow arrowheads) extended into the ependymal cell layer (indicated by PI-stained cells) adjacent to the 3V. (G–J) Statistical results showing the differences of somatic volume (G) (P = 0.0119), Herring body number (H) (P = 0.0005), total Herring body volume per cell (I) (P = 0.0003), and the average volume of the Herring bodies (J) (P = 0.0047) between neurons in the PaAp and Pe. (K) Correlation between somatic volume and Herring-body-positive number of neurons in the PaAp. Spearman’s correlation coefficient was r = −0.4519, P = 0.0455. Each point in the scatterplot represents a single cell. (L and M) There were no significant difference in fibrous features (L, total fibrous length; M, total number of fibrous branches) of neurons between PaAp and Pe. ns: not significant. (N) Comparison of primary fibrous branches and the proportions of bipolar or multipolar cells in the PaAp and Pe.

In the Pe, the reconstructed somata were located in the lower part of the PaAp and around the 3V (Fig. 5 C, purple bodies; Fig. S4E indicated by red circles). These cells sent out fibers and one or two of them extended close to the 3V (Fig. 5C and D, red line). There were large Herring bodies on the fibers (Fig. 5D, grey reconstituted bodies; Fig. 5F, indicated by yellow arrowheads) and they terminated in the ependymal cell layer adjacent to the 3V (Fig. 5F). Most of the reconstructed cells within the Pe were bipolar neurons (Fig. 5D, e and f), and the fibers extending downward to the 3V had more Herring bodies (Fig. 5D, f and g).

Next, we compared the somatic and fiber parameters of the neurons in the PaAP and Pe and found that the average volume of the soma in the Pe (951.2±108.3 μm^3^) was larger than that in the PaAP (640.1±60.4 μm^3^) (Fig. 5G, P=0.0119), and the number of Herring bodies in the Pe (19.4±2.5) was significantly greater than that in the PaAP (7.9±1.6) (Fig. 5H, P=0.0005). The total Herring-body volume (2678±652.3 μm^3^) (Fig. 5I, P=0.0003) per cell and the average volume of Herring bodies (153.4±43.1 μm^3^) (Fig. 5J, P=0.0047) in the Pe were significantly larger than those (468.2±79.2 μm^3^ and 47.7±5.3 μm^3^) in the PaAP. We also found that there was a negative correlation between the number of Herring bodies and the soma volume in the PaAP (r=−0.4519, P=0.0455) (Fig. 5K). There was no significant difference in the total fibrous length (PaAP: 455.4±24.1 μm, Pe: 537.5±57.6 μm) (Fig. 5L) or the total number of fibrous branches (PaAP: 5.4±0.9, Pe: 6.3±1.6) (Fig. 5M) of neurons in the PaAP and Pe. We found that most (75% in the PaAP and 70% in the Pe) (Fig. 5N, right half, indicated in the pie chart) of the reconstructed neurons were bipolar neurons, which were characterized by the number of primary fibrous branches (Fig. 5N, left half).

Collectively, these 3D reconstructions of hypothalamic CRH neurons may be indicative of the transport and storage of CRH peptides in hypothalamic neurons, as well as the possible their release sites, such as the third ventricle. These findings provide a structural basis for further elucidating the neural circuits and functions of CRH neurons.

## Discussion

During the last few decades, transgenic rodent models have become powerful tools for studying the distribution (Chen et al., 2015; Kono et al., 2017; Wamsteeker Cusulin et al., 2013) and function (Kolber et al., 2010; Lu et al., 2008; Regev et al., 2012; Stenzel-Poore et al., 1994) of CRH neurons in the brain. In order to probe the morphological characteristics of CRH neurons at single-cell resolution, we combined genetic labeling (using transgenic mouse lines) with the fMOST platform to generate high-resolution imaging datasets, with which we characterized the morphologies of distinct CRH neurons distributed in various brain regions throughout the whole mouse brain.

The robust native fluorescence of each of these reporter mouse lines enabled direct visualization of fine dendritic and axonal structures of labeled neurons, which is demonstrated to be useful for mapping neuronal circuitry, as well as imaging and tracking specific cell populations (Madisen et al., 2010; Taniguchi et al., 2011; Walker et al., 2019). We compared the distribution patterns of fluorescent-labeled CRH neurons in three reporter mouse lines. Notably, the fluorescent fusion protein, CHR2-EYFP, is membrane-bound and is therefore distributed along the plasma membrane of neuronal processes within CRH-IRES-Cre;Ai32 mice (Dedic et al., 2018; Park et al., 2018), which enables a clear visualization of the entire neuronal morphology. Therefore, we utilized CRH-IRES-Cre;Ai32 mice for whole-brain imaging and reconstructions. Interestingly, a large number of EYFP-labeled cortical pyramidal neurons was also observed in adult mice (which has not been reported previously from the onset age of postnatal day 21) (Fig. S3).

Next, we focused our analyses of reconstructed neurons on several stress-related regions, including the mPFC, hypothalamus, amygdala, BST, and hippocampus. For example, it has been reported that local CRH-synthesizing neurons are prominent in the PFC (Charlton et al., 1987; De Souza et al., 1985; Lewis et al., 1989; Merchenthaler et al., 1984; Swanson et al., 1983) and may modulate the activities of pyramidal neurons (Gallopin et al., 2006). However, until now, the complete morphologies of CRH neurons in the mPFC have rarely been reported. Here, we reconstructed fluorescent-labeled CRH neurons in the cortical column in the PrL within mPFC across layers 1–4 (Fig. 4A). Importantly, we classified different neuronal types by their soma depths and arborization patterns (example listed in Fig. 4B). For the first time, we showed the distribution of CRH neurons with different morphological types in different cortical layers. We found there were dense dendrites (Fig. 4A) with dendritic swellings (Fig. S3, b) in layer 1 (the soma of which were located in layer 2/3 or layer 4), and that fibers extended to the surface of the cortex. According to the layer-specific dendritic locations and their different projection targets, several types of putative connection patterns between CRH neurons were identified in the cortex (listed in Fig. 4F–H). Such a diverse dendritic connection pattern of cortical CRH neurons may reflect differential innervation of downstream output targets (each amplified subfigure shown in Fig. 4F–H). Therefore, by characterizing their somatic locations and unique respective local dendritic morphologies of CRH neurons, our present study not only increases our current understanding of the distribution of CRH neurons, but also enables future studies to further elaborate upon cell-specific classifications. Taken together, these findings may help elaborate future functional studies of morphologically diverse CRH neurons in the PFC.

Importantly, CRH functions as a neuropeptide hormone produced in neuroendocrine neurons in the PVN and regulates the synthesis and secretion of glucocorticoids from the adrenal glands through the action of adrenocorticotropic hormone. For the first time, we reconstructed the intact morphologies of CRH neurons and their neurite varicosities (Herring bodies) located within fibers in the PaAp (Fig.5 A–B) and Pe CRH neurons (Fig. 5C–D). In the PaAp, most fibers with Herring bodies projected rostrally (Fig. 5B), while in the Pe, the ends of the fibers extended to the third ventricle (Fig. 5C–D and F). Since Herring bodies mainly function to store and secrete CRH (Dellmann and Rodriguez, 1970; Vazquez and Amat, 1978; Yamazaki et al., 1981), we speculate that these endocrine CRH neurons are different from those that project to the median eminence, and that CRH may be also secreted to other areas of the hypothalamus or cerebrospinal fluid to participate in its regulatory functions. We further found a negative correlation between somatic volume and Herring body number in the PaAp (Fig. 5K). Thus, our reconstructed morphological characteristics of Herring bodies (located in the fibers of CRH neurons) may facilitate future classifications (according to different fiber orientations and Herring-body distribution patterns) of hypothalamic CRH neurons and advance our understanding of their potentially diverse functions.

The reconstructed CRH neurons in different brain regions showed diverse distribution patterns and morphologies (Fig. 3A–H). We found that some neurons shared a common bipolar shape across various brain regions (Fig. 3I), especially in the hypothalamus (75% in the PaAp, 100% in SCN) and cortex. Rho *et al.* reported that parvocellular CRH neuroendocrine neurons typically have two relatively thick primary dendrites that extend from opposite sides of the soma in a bipolar arrangement and branch once (Rho and Swanson, 1989); and furthermore, bipolar cells are commonly found in the cortex (Cahusac et al., 1998; Delville et al., 1992; Yan et al., 1998). Thus, our present study in CRH-reporter mice is consistent with these previous studies and is the first to describe the specific neural structures of these CRH neurons, such as the different types of connections between CRH neurons in the mPFC, as well as the intact morphologies of Herring bodies in hypothalamic CRH neurons. Such simplified branching properties for these CRH neurons in the hypothalamus may be conducive to their endocrine functions. Among all the reconstructed CRH neurons across different brain regions, their somata had different sizes (Fig. 3J), with somata in the LSD, VMPO, and hippocampus being larger than those in the PaAp, Pe, and SCN. Also, we found differential dendritic branch complexities across these regions. For example, CRH neurons in the mPFC, LSD, and hippocampus all exhibited more complex dendritic morphologies compared to those in the PaAP, Pe, and SCN (Fig. 3K–L). Hupalo *et al.* demonstrated that chemogenetic activation of caudal but not rostral dmPFC CRH neurons potently impaired working memory, whereas inhibition of these neurons improved working memory (Hupalo et al., 2019b). In addition, CRH acts in the medial septum to impair spatial memory (Hupalo et al., 2019a), and acts in the BNST to participate in stress-induced maladaptive behaviors (Hu et al., 2020). However, the functions of CRH in the OB, SCN, and VMPO remain unclear. Therefore, our current study may provide a detailed morphological basis for future functional-based studies for CRH neurons in these different brain regions. Interestingly, the PaAP, Pe, and SCN all belong to the hypothalamus, and CRH neurons in these regions had smaller somata and exhibited more prevalent bipolar branching patterns compared to other brain regions. In the PVN, CRH neurons have been identified as parvocellular cells (Miklos and Kovacs, 2002; Rho and Swanson, 1989). We found that the mean somatic volume of CRH neurons in the Pe was larger than those in the PaAP (Fig. 5G). Hence, we speculate that these data may be indicative of two different types of CRH endocrine neurons within the hypothalamus. Collectively, our quantitative analysis of these reconstructions demonstrates a region-specific diversity of CRH neurons in terms of both somatic size and branching complexity.

In summary, in the present study, we generated high-resolution imaging datasets to characterize, at single-cell resolution, the fine morphologies of CRH neurons distributed in diverse brain regions. Such region-specific reconstructions of intact morphologies of CRH neurons may help in further elucidating both CRH-mediated physiological functions in various brain circuits and the associations of their dysfunction with various neuropathological diseases.

## Materials and Methods

### Animals

CRH-IRES-Cre (B6(Cg)-Crh^tm1(cre)Zjh^/J; stock number: 012704), Ai6 (B6.Cg-Gt(ROSA)26Sor^tm6(CAG-ZsGreen1)Hze^/J; stock number: 007906), Ai14 (B6.Cg-Gt(ROSA)26Sor^tm14(CAG-TdTomato)Hze^/J; stock number: 007914) and Ai32 (B6;Cg-Gt(ROSA)26Sor^tm32(CAG-COP4*H134R/EYFP)Hze^/J; stock number: 012569) mice have been described previously(Madisen et al., 2012; Madisen et al., 2010; Taniguchi et al., 2011). CRH-IRES-Cre, Ai6 and Ai32 mice were purchased from the Jackson Laboratory. Ai14 mice were obtained from the laboratory of Minmin Luo (NIBS, China). All of the mice were bred onto a C57BL/6J genetic background. CRH-IRES-Cre;Ai6, CRH-IRES-Cre;Ai14 and CRH-IRES-Cre;Ai32 mice were derived from crosses of the CRH-IRES-Cre and Ai6, Ai14 and Ai32 genotypes, respectively. Male mice at 12 weeks of age were used for experiment. CRH-IRES-Cre;Ai32 male mice were used for fMOST imaging and neuron reconstructions. The mice were housed on a 12-hour light/dark cycle with food and water ad libitum. All the animal experiments were performed according to the procedures approved by the Institutional Animal Ethics Committee of University of Science and Technology of China.

### Histology

All histological procedures have been previously described(Gong et al., 2016; Xiong et al., 2014; Yang et al., 2013). Briefly, for whole-brain imaging, the mice were anaesthetized and perfused with 0.01M PBS (Sigma-Aldrich Inc., St. Louis, USA), followed by 4% paraformaldehyde (PFA) and 2.5% sucrose in 0.01M PBS. The brains were excised and post-fixed in 4% PFA for 24 h. After fixation, each intact brain was rinsed overnight at 4°C in 0.01M PBS and subsequently dehydrated in a graded ethanol series. Then, the brains were impregnated with Glycol Methacrylate (GMA, Ted Pella Inc., Redding, CA) and embedded in a vacuum oven.

For immunofluorescence and visualizing fluorescent labeled neurons in three kinds of mice lines, the fixed brains were embedded by agarose and consecutive 50 or 100 μm coronal sections were collected using a vibration microtome (Leica VT1200S, Germany). For CRH immunofluorescence, 50 μm sections were washed in PBS three times (10 min each time) and permeabilized with 0.3% Triton X-100 for 30 min, followed by 5% normal donkey serum blocking solution at room temperature for 1 h. Sections were then incubated with rabbit anti-CRF (1:2,000, Bachem, T4037) primary antibody in PBS containing 0.3% Triton X-100 overnight or for 36 h at 4°C. After washing in PBS, sections were incubated with Alexa Fluor 594-conjugated donkey anti-rabbit secondary antibody (1:200, Jackson Immuno Research, 711-585-152) diluted in 0.1% Triton X-100 in PBS at room temperature for 2 h. After washing in PBS, sections were mounted on slides with antifade mounting medium (Vector Laboratories, Inc, H-1000) and stored at 4°C. For visualizing fluorescent labeled neurons, 100 μm sections were washed in PBS and mounted on slides. All images were photographed using LSM 880 (Zeiss, Germany) confocal microscope.

### Whole-brain imaging

Whole-brain imaging was performed by the fMOST system(Gong et al., 2016). Briefly, the immersed samples were fixed on the imaging plane, a WVT system automatically performed the sectioning and imaging to complete the brain-wide data acquisition. We acquired the data sets after sectioning at a 1 um thickness and imaging at a 0.32 × 0.32 ×1 um voxel size. To enhance the in-focus EYFP signal, we added Na_2_CO_3_ into the water bath. Most of the EYFP molecules were preserved in a nonfluorescent state, rather than directly damaged, through chromophore protonation during the resin-embedding procedure. These fluorescent signals were chemically recovered to the fluorescent state using 0.05 M Na_2_CO_3_ during imaging. For the CRH-IRIS-Cre;Ai32 samples, real-time PI staining was performed.

### Image preprocessing

The raw data acquired by the fMOST system needed image preprocessing for mosaic stitching and illumination correction. This process has been described before(Gong et al., 2016). Briefly, the mosaics of each coronal section were stitched to obtain an entire section based on accurate spatial orientation and adjacent overlap. Lateral illumination correction was performed section by section. Image preprocessing was implemented in C++ and optimized in parallel using the Intel MPI Library (v.3.2.2.006, Intel). The whole data sets were executed on a computing server (72 cores, 2GHz per core) within 6h. All full coronal sections were saved at an 8-bit depth in LZW compression TIFF format after image preprocessing.

### Visualization and reconstruction

We visualized the data set using Amira software (v.5.2.2, FEI) and Imaris software (v.9.2.1, bitplane, Switzerland) to generate the figures and movies. To process the TB-sized data on a single workstation, we transformed the data format from TIFF to the native LDA type using Amira. The visualization process included extracting the data in range of interest, sampling or interpolation, reslicing the images, identifying the maximum intensity projection, volume and surface rendering, and generating the Movies using the main module of Amira. The segmentation editor module of Amira was utilized for the manual outline segmentation of the third ventricle, somata and Herring bodies. We applied the filament editor module of Amira to trace the morphology of EYFP-labeled neurons in 3D by human-machine interaction. The reconstructed neurons were checked back-to-back by three persons. The tracing results with original position information were saved in SWC format and the results of the analyses were generated by L-Measure software (v.5.3).

### Statistics

All statistical graphs were generated using Graphpad Prism v.6.01. Two-tailed Student’s t-test and one-way ANOVA followed by Tukey’s post hoc tests were also performed using Graphpad Prism v.6.01 and SPSS (IBM SPSS Statistics 23). The confidence level was set to 0.05 (P value) and all results were presented as the means ± S.E.M.

## Acknowledgments

We thank Dr. Minmin Luo (NIBS) for providing the Ai14 mice; Xingning Li, Miao Ren, Beng Long, Jie Peng, Yuxin Li (HUST), Hui Fang, Xinya Qin, and Penghao Luo (USTC) for help with the experiment.

