## Supplementary Information for "Visualizing dendritic characteristics of corticotropin-releasing hormone neurons at single-cell resolution in the whole mouse brain"

* Hui Gong or Jiang-Ning Zhou

**This PDF file includes:**

Figures S1 to S4

Tables S1 to S2

Legends for Movies S1 to S3

**Other supplementary materials for this manuscript include the following:**

Movies S1 to S3

### Supplementary Figures:

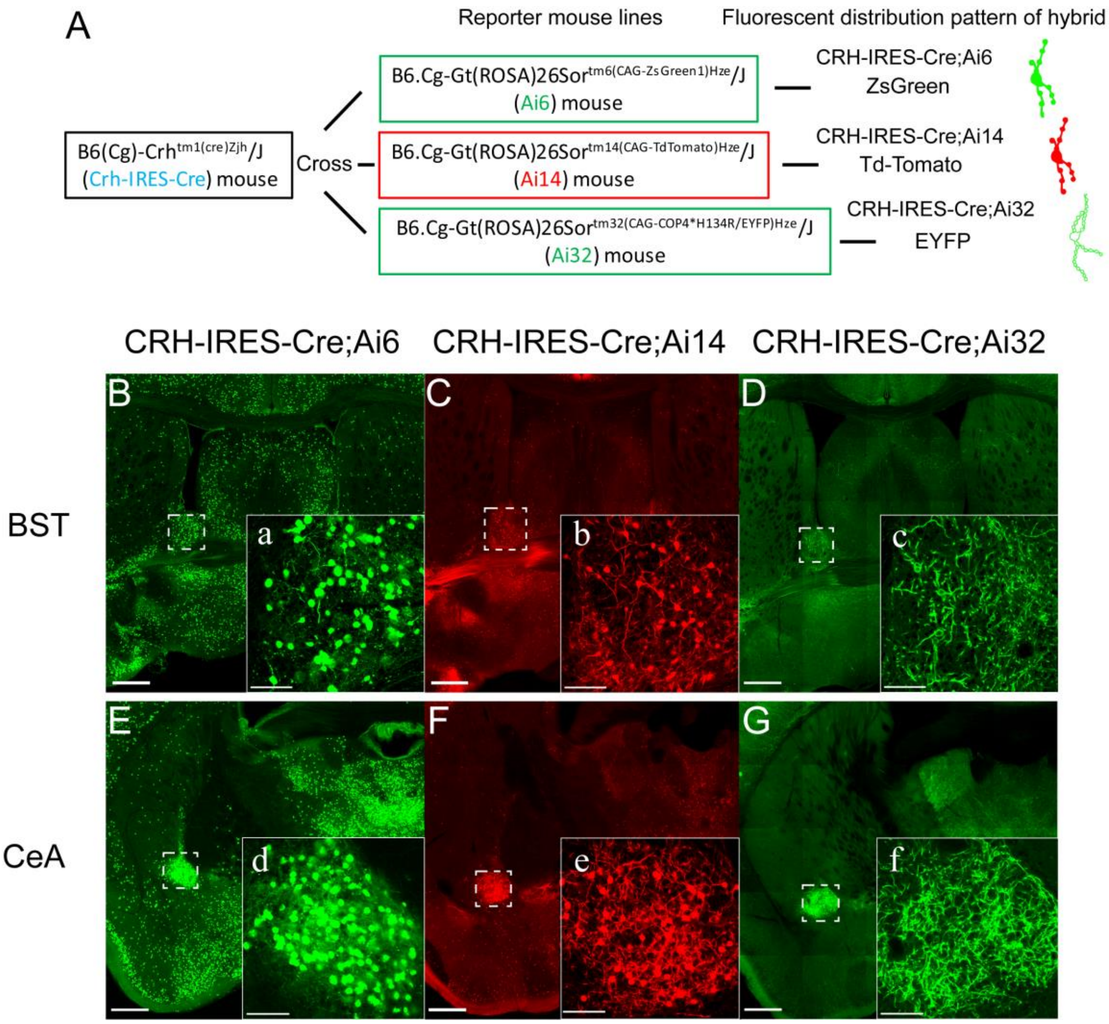


**Fig. S1. Generation of transgenic mouse lines and morphological features of CRH neurons in the BST and CeA.**

(A) Schematic diagram illustrating the generation of three transgenic mouse lines. Breeding CRH-IRIS-Cre mice with Ai6, Ai14, and Ai32 mice yielded CRH-IRIS-Cre:Ai6, CRH-IRIS-Cre:Ai14, and CRH-IRIS-Cre:Ai32 mice, in which ZsGreen1, Td-Tomato, and EYFP, respectively, were expressed specifically in cell bodies or fibers of CRH-positive neurons. (B–D) Distributions and morphologies of fluorescent-labeled CRH neurons in the BNST of the three mouse lines. a–c: Magnified images from the dotted boxes in B, C, and D, respectively. (E–G) Distributions and morphologies of fluorescent-labeled CRH neurons in the CeA of the three mouse lines. d–f: Magnified images from the dotted boxes in E, F and G, respectively. Scale bars = 500 μm and 100 μm for the inserts.

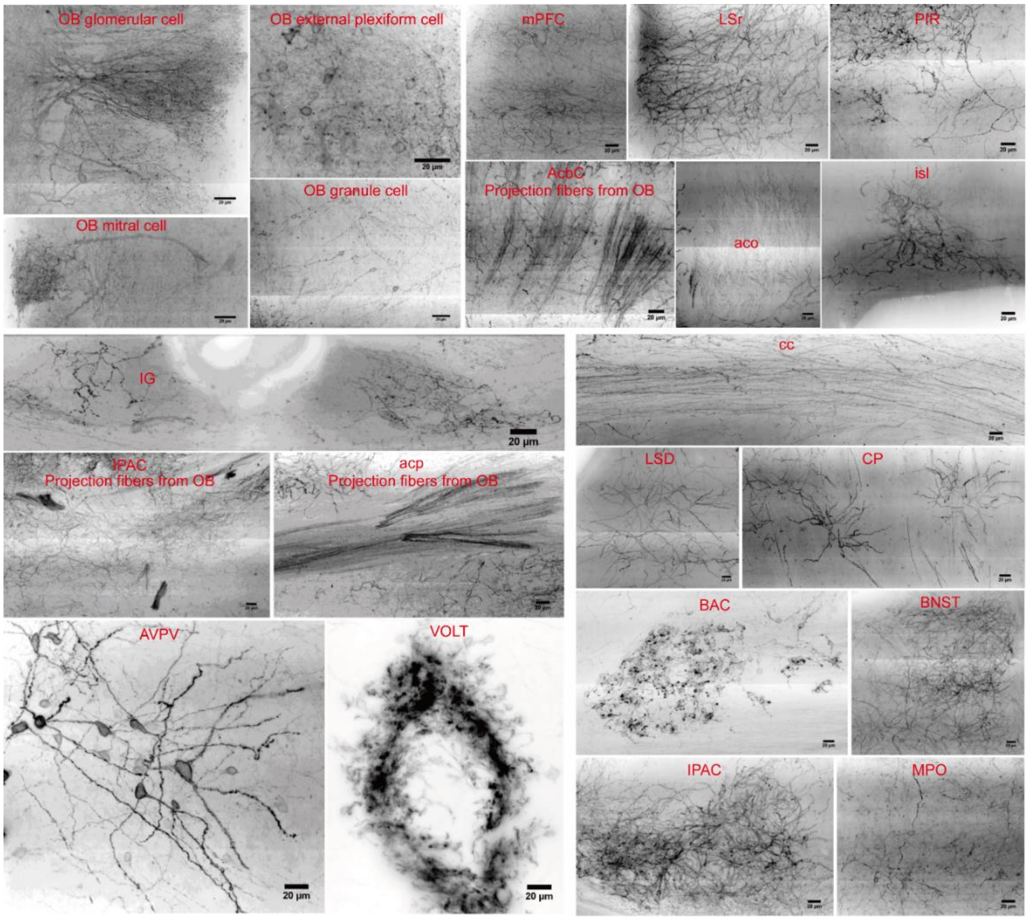
OB glomerular cell
OB external plexiform cell
mPFC
LSr
PIR
OB mitral cell
OB granule cell
AcbC
Projection fibers from OB
aco
isl
IG
cc
IPAC
Projection fibers from OB
acp
Projection fibers from OB
LSD
CP
BAC
BNST
AVPV
VOLT
IPAC
MPO
20 μm

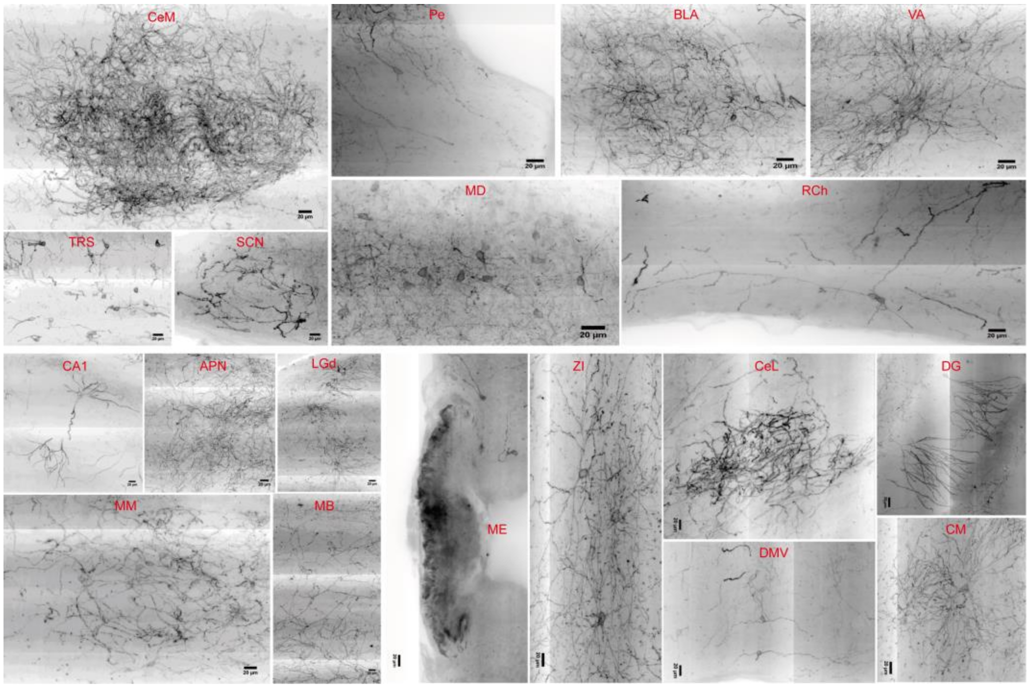
CeM
Pe
BLA
VA
TRS
SCN
MD
RCh
CA1
APN
LGd
ME
ZI
CeL
DG
MM
MB
DMV
CM
20 μm

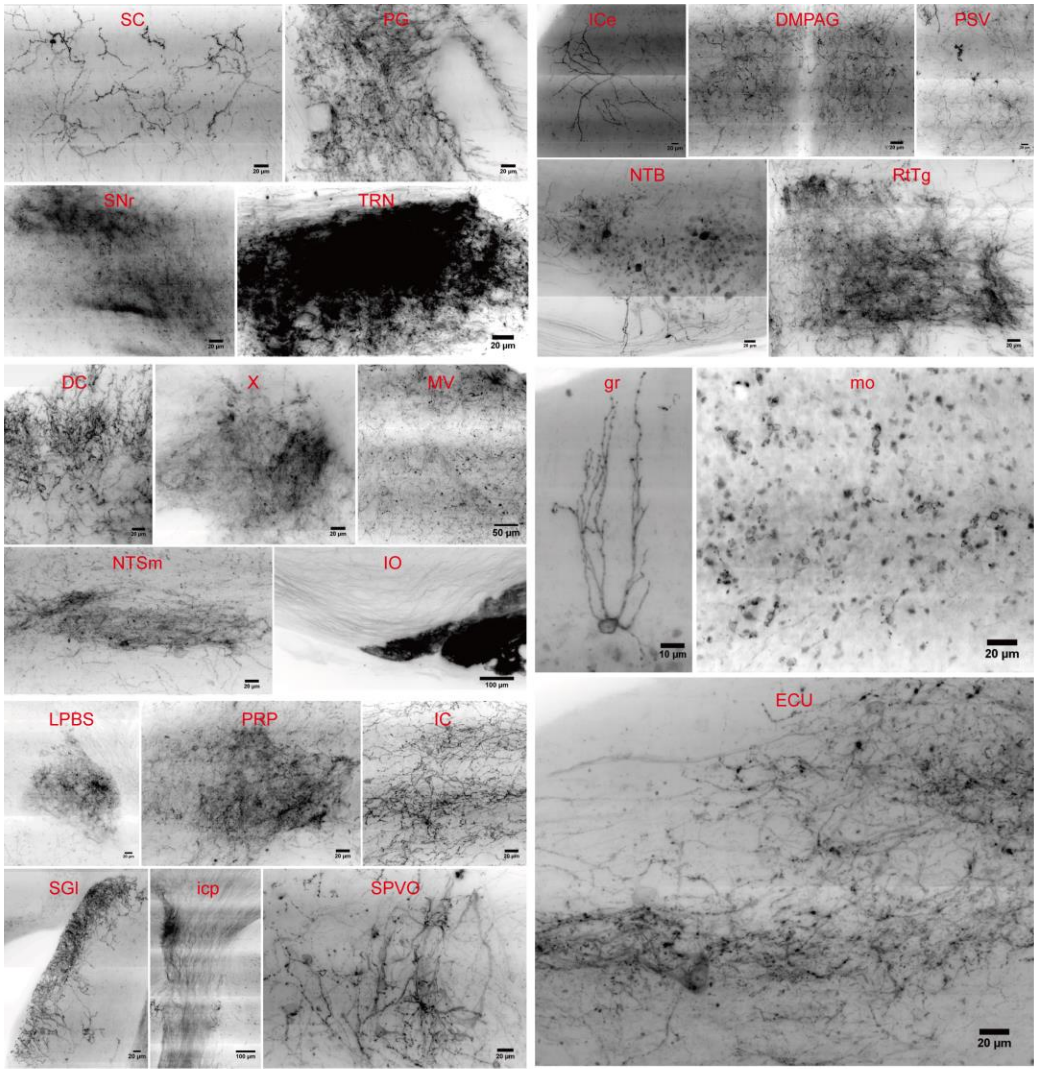


**Fig. S2. High-resolution images showing diverse morphologies of CRH neurons throughout the brains of CRH-IRIS-Cre;Ai32 mice.**

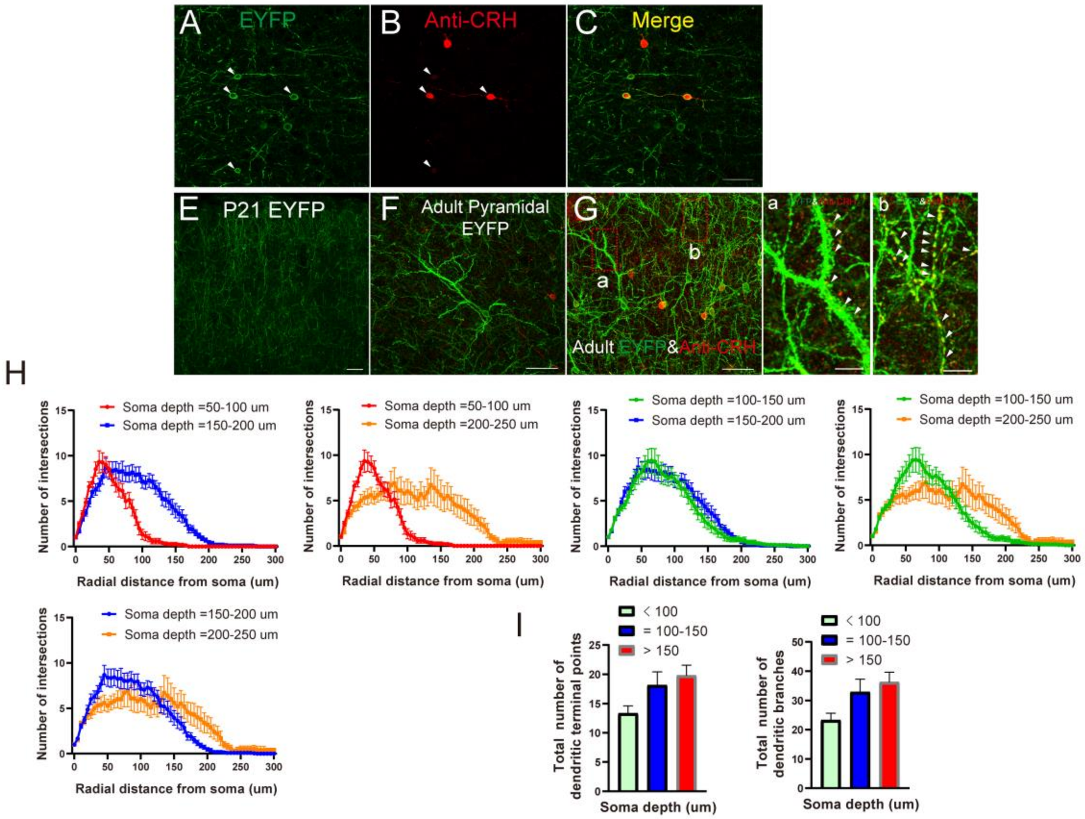


**Fig. S3. Expression specificity and dendritic analysis in the mPFC of CRH-IRIS-Cre;Ai32 mice.**

(A–C) Immunofluorescent staining showing that EYFP and CRH were co-labeled (indicated by arrowheads) in the mPFC. (E–F) EYFP-labeled pyramidal CRH neurons were visible in adult mice in the mPFC. (G) EYFP-labeled pyramidal neurons included the apical dendrites (red dotted box a) and dendritic spines (indicated by arrowheads in magnified image a) but were not co-labeled with CRH antibodies, while the apical dendrites (red dotted box b) and dendritic swellings (indicated by arrowheads in magnified image b) were CRH-positive. Scale bars: A–G: 50 µm, a–b: 10 µm. (H) Sholl analysis of dendrites of neurons with different somatic depths illustrating changes in the mean number of intersections with increasing radial distance from the soma. (I) The total number of dendritic terminal points and total number of dendritic branches were not significantly different among neurons with somatic depths < 100 µm, soma depths = 100–150 µm, and somatic depths > 150 µm.

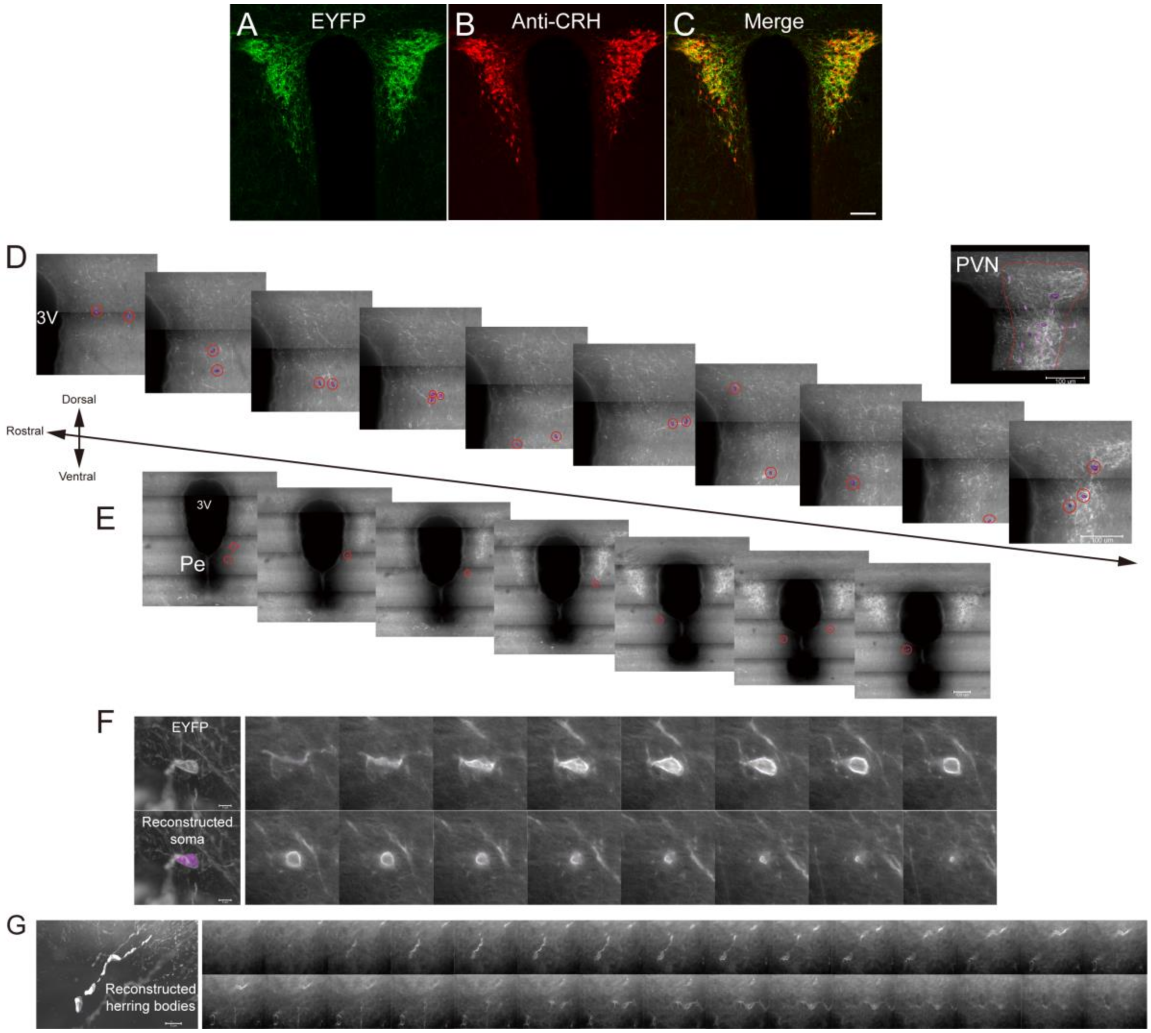


**Fig. S4. Expression specificity in PaAp, somatic locations of reconstructed neurons in the PaAp and Pe, and examples of the reconstructions of somata and Herring bodies in CRH-IRIS-Cre;Ai32 mice.**

(A–C) Immunofluorescent staining showing that EYFP and CRH were co-labeled in the PaAp. Scale bar: 100 μm (D) The somatic locations (indicated by red circles) of each reconstructed neuron in the PaAp. (E) The somatic locations (indicated by red circles) of each reconstructed neuron in the Pe. (F–G) 3D images and consecutive 1-μm sections showing the reconstruction process of somata (F) and Herring bodies (G).

| Brain region | Soma volume (x10³μm³) | Total dendritic length (mm) | Number of dendritic branches |
|---|---|---|---|
| mPFC | 0.40±0.02 | 1.37±0.19 | 44.21±4.01 |
| LSD | 1.67±0.12 | 1.90±0.22 | 27.71±2.99 |
| BST | 1.02±0.14 | 1.29±0.13 | 16.71±3.62 |
| VMPO | 1.17±0.23 | 1.07±0.13 | 15.20±3.67 |
| Hip | 0.93±0.10 | 1.64±0.10 | 24.75±2.94 |
| CeA | 1.21±0.06 | 1.38±0.38 | 39.50±7.97 |
| PaAp | 0.64±0.06 | 0.46±0.02 | 5.40±0.86 |
| Pe | 0.95±0.11 | 0.54±0.06 | 6.30±1.56 |
| SCN | 0.64±0.06 | 0.56±0.05 | 6.20±0.68 |

**Table S1. Parameters of somatic volume, total dendritic length, and the number of dendritic branches of the reconstructed neurons in several brain regions.**

**Table S2. Abbreviation for brain area.**

| | | | |
|---|---|---|---|
| 3V | third ventricle | IL | infralimbic cortex |
| AcbSh | accumbens nucleus,shell | IO | inferior olive |
| acp | anterior commissure, posterior | IPAC | interstitial nucleus of the posterior limb of the anterior commissure |
| AD | anterodorsal thalamic nucleus | KF | Kolliker-Fuse nucleus |
| AM | anteromedial thalamic nucleus | LD | laterodorsal thalamic nucleus |
| APTD | anterior pretectal nucleus, dorsal part | LH | lateral hypothalamic area |
| Au1 | primary auditory cortex | LS | lateral septal nucleus |
| AuD | secondary auditory cortex, dorsal area | LSD | lateral septal nucleus, dorsal part |
| BAC | bed nucleus of the anterior commissure | M1 | primary motor cortex |
| BST | bed nucleus of the stria terminalis | M2 | secondary motor cortex |
| CA1 | field CA1 of hippocampus | MD | mediodorsal thalamic nucleus |
| cc | corpus callosum | ME | median eminence |
| CeA | central amygdaloid nucleus | Mi | mitral cell layer |
| Cg1 | cingulate cortex, area 1 | MGV | medial geniculate nucleus, ventral part |
| CPu | caudate putamen | MM | medial mammillary nucleus, medial part |
| DC | dorsal cochlear nucleus | MnR | median raphe nucleus |
| DG | dentate gyrus | MVe | medial vestibular nucleus |
| DLG | dorsal lateral geniculate nucleus | OB | olfactory bulb |
| DM | dorsomedial hypothalamic nucleus | PaAp | paraventricular hypothalamic nucleus, anterior parvicellular part |
| DTT | dorsal tenia tecta | Pe | periventricular hypothalamic nucleus |
| ECu | external cuneate nucleus | PF | parafascicular thalamic nucleus |
| EPl | external plexiform layer | Pir | piriform cortex |
| Gl | glomerular layer | Pn | pontine nuclei |
| GrO | granule cell layer | Po | posterior thalamic nuclear group |
| Hip | hippocampus | PR: | prerubral field |

| | | | |
|---|---|---|---|
| IC | inferior colliculus | Pr5 | principal sensory trigeminal nucleus |
| icp | inferior cerebellar peduncle | PrL | prelimbic cortex |
| PVN | paraventricular hypothalamic nucleus | V1 | primary visual cortex |
| RSG | retrosplenial granular cortex | V2 | secondary visual cortex |
| RtTg: | reticulotegmental nucleus of the pons | VCA | ventral cochlear nucleus, anterior part |
| S1 | primary somatosensory cortex | VCP | ventral cochlear nucleus, posterior part |
| S2 | secondary somatosensory cortex | VLG | ventral lateral geniculate nucleus |
| SCN | suprachiasmatic nucleus | VLTg | ventrolateral tegmental area |
| SNC | substantia nigra, compact part | VMPO | ventromedial preoptic nucleus |
| SuG | superficial gray layer of the superior colliculus | VOLT | vascular organ of the lamina terminalis |
| TS | triangular septal nucleus | X | nucleus X |

**Movie S1. Movie of serial sections showing the fiber projections from OB.**

**Movie S2. Movie of serial sections showing the fiber projections from IO.**

**Movie S3. 3 D movie showing the type II and type III connections of CRH neurons in the mPFC.**
